# Innovative multidimensional models in a high-throughput-format for different cell types of endocrine origin

**DOI:** 10.1101/2022.01.10.475686

**Authors:** Stefan Bornstein, Igor Shapiro, Maria Malyukov, Richard Züllig, Edlira Luca, Evgeny Gelfgat, Felix Beuschlein, Svenja Nölting, Alfredo Berruti, Sandra Sigala, Mirko Peitzsch, Charlotte Steenblock, Barbara Ludwig, Patrick Kugelmeier, Constanze Hantel

**Affiliations:** Department of Endocrinology, Diabetology and Clinical Nutrition, University Hospital Zurich (USZ) and University of Zurich (UZH), Zurich, Switzerland; Kugelmeiers AG, Erlenbach, Switzerland; Oncology Unit, Department of Medical and Surgical Specialties, Radiological Sciences, and Public Health, University of Brescia at ASST Spedali Civili di Brescia, 25123 Brescia, Italy; Endocrine Research Unit, Medizinische Klinik und Poliklinik IV, Klinikum der Universität München, 80336 Munich, Germany; Department of Medicine IV, University Hospital, LMU Munich, Ziemssenstraße 1, 80336 München, Germany; Section of Pharmacology, Department of Molecular and Translational Medicine, University of Brescia, 25123 Brescia, Italy; Medizinische Klinik und Poliklinik III, University Hospital Carl Gustav Carus Dresden, Dresden, Germany; Institute of Clinical Chemistry and Laboratory Medicine, University Hospital Carl Gustav Carus at Technische Universität Dresden, Dresden, Germany

**Keywords:** 3D-models, pseudoislets, adrenal, adrenocortical carcinoma, adrenocortical cell lines, NCI-H295, MUC-1, spheroids, organoids, cell-replacement therapies, cancer

## Abstract

The adrenal gland provides an important function by integrating neuronal, immune, vascular, metabolic and endocrine signals under a common organ capsule. It is the central organ of the stress response system and has been implicated in numerous stress-related disorders. While for other diseases, regeneration of healthy organ tissue has been aimed at such approaches are lacking for endocrine diseases - with the exception of type-I-diabetes. Moreover, tumor formation is very common, however, appropriate high-throughput applications reflecting the high heterogeneity and furthermore relevant 3D-structures in vitro are still widely lacking. Recently, we have initiated the development of standardized multidimensional models of a variety of endocrine cell/tissue sources in a new multiwell-format. Firstly, we confirmed common applicability for pancreatic pseudo-islets. Next, we translated applicability for spheroid establishment to adrenocortical cell lines as well as patient material to establish spheroids from malignant, but also benign adrenal tumors. We aimed furthermore at the development of bovine derived adrenal organoids and were able to establish steroidogenic active organoids containing both, cells of cortical and medullary origin. Overall, we hope to open new avenues for basic research, endocrine cancer and adrenal tissue-replacement-therapies as we demonstrate potential for innovative mechanistic insights and personalized medicine in endocrine (tumor)-biology.

## Introduction

Spheroids and organoids have become increasingly popular to study organogenesis and have opened avenues to drug discovery and personalized medicine. In cancer biology, multiple studies have demonstrated the genetic and histological comparability between cultured tumor spheroids and the original tumor, where the 3D cultures successfully recapitulate the phenotypic profile of the tumor growing in vivo (de Witte et al., 2020; Driehuis et al., 2019; Gao et al., 2014; Sachs et al., 2018; Tiriac, Plenker, Baker, & Tuveson, 2019). As such, 3D cultures are being used to better understand the response to various drugs and therapeutics, involving various additional factors such as tumor microenvironment and multi-layer structures and, thereby, defining individualized treatments for solid tumors (B. Pinto, Henriques, Silva, & Bousbaa, 2020).

However, the future of these multifaceted structures in the endocrine field goes far beyond and includes transplantation and organ replacement in humans in certain pathologies, such as type I diabetes, adrenal insufficiency or refractory Cushing’s disease (Bornstein et al., 2020; Dong, Ji, & Li, 2019; Lanzoni & Ricordi, 2021). Moreover, in islet research, which lacks different suitable cell lines, intact islets can be isolated from multiple donor specimens and maintained in vitro. However, these are impermeable to genetic manipulations since viral transduction only reaches the outermost cells, while the inner regions of the islets are not affected (Friedlander, Nguyen, Kim, & Bevacqua, 2021). Dispersion of the islets and their subsequent re-aggregation into spheroids allows for the manipulation of all cells and the consecutive study within a multi-layer structure.

Similarly, in vitro models of adrenal origin have suffered from a lack of suitable human cell lines that resemble functional properties of benign and malignant tumors. Fortunately, this gap could recently been narrowed with cell models that are now applicable for a spectrum of research applications (Abate et al., 2020; Fragni et al., 2019; Hantel et al., 2016; Hasanovic et al., 2018; Landwehr et al., 2021; Liang et al., 2020; E. M. Pinto, Kiseljak-Vassiliades, & Hantel, 2019; Rossini et al., 2021; Siebert et al., 2019; Warde et al., 2020). However, there is still potential for improvement by translating adrenocortical cells towards 3D culture and appropriate personalized approaches in vitro. Moreover, for the diverse types of adrenocortical adenoma and paraganglioma/pheochromocytoma, appropriate in vitro models of human origin are still urgently needed (Bayley & Devilee, 2020).

Overall, there are manifold preclinical and clinical implications for tumor spheroid and organoids derived from human endocrine tissue origins. Thus, innovative high throughput applications for 3D models of endocrine origin open new avenues for endocrine cell replacement therapies and other personalized therapeutic approaches.

## Results

### Plate design

The Sphericalplate 5D is a 24-well laboratory plate with specially designed and patented microwells (Figure 1A).

**Figure 1:**
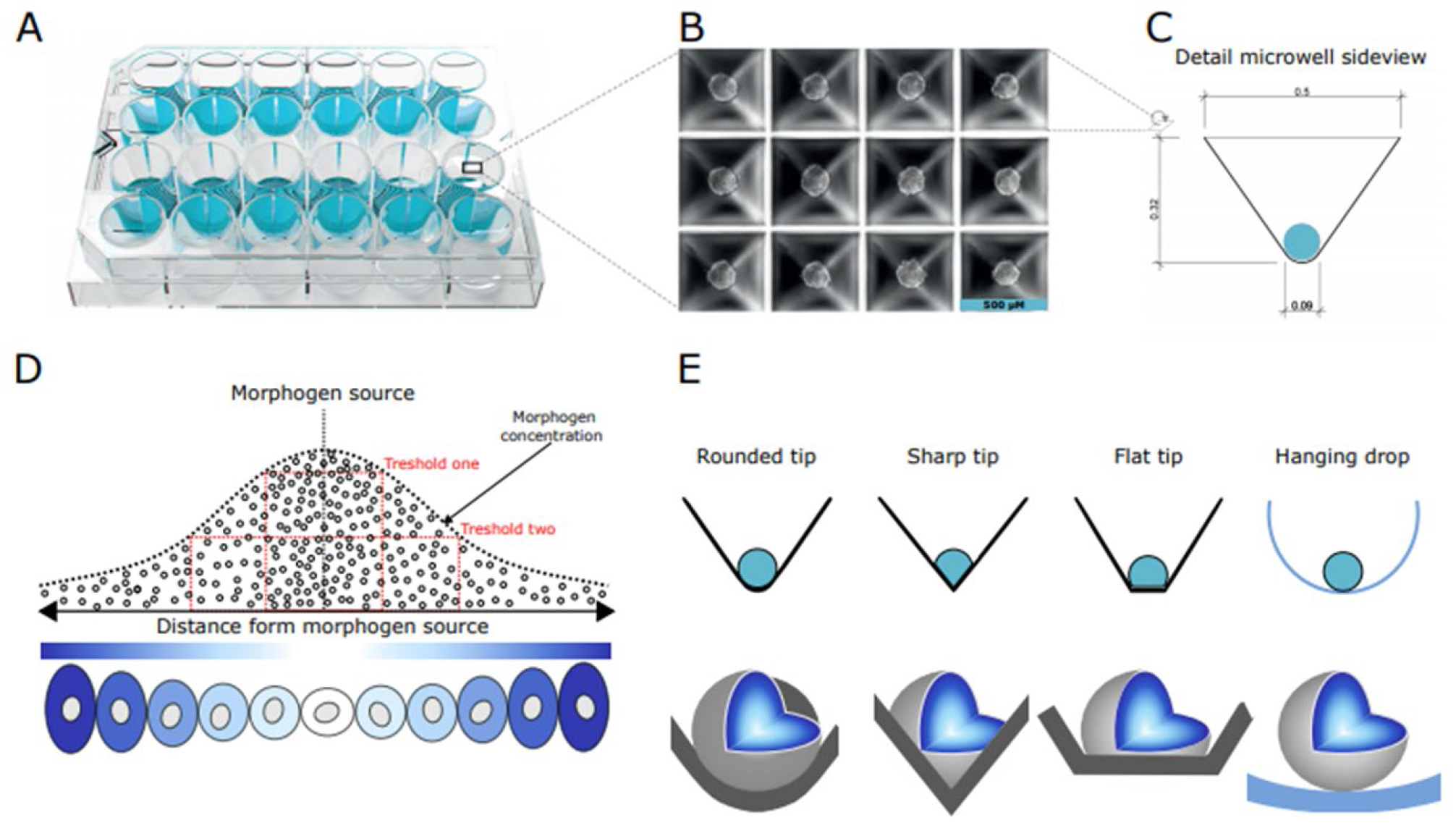
Layout of the Sphericalplate 5D highlighting the 12 working wells (A). Image of spheroids formed in the microwells within one of the wells of the plate (B). Shape and measurements of the microwell (C). Concentration gradient of morphogen (D,(Huizar, Soundarrajan, Paravitorghabeh, & Zartman, 2020)). Advantage of rounded-tip design of microwell on spheroid formation (E-F).

Each well is set up to allow the easy production of 750 uniform and size-controlled islets or other cell spheroids with only one pipet hub of a respective cell suspension (Figure 1B). The spherical geometry with square top openings ensures that all cells participate in spheroid formation (Figure 1C). The top square openings then merge into rounded tips to ensure the creation of uniform spheroids with cellular functionality and viability. This is achieved serving fundamental biological principles like free energy minimisation with optimal tip geometry and special wall angles supporting spheroids but not limiting them. In addition, a patented medical grade ultra-low attachment coating ensures minimal surface interaction. Thus, compared to other available plate types, there is no need for rinsing solutions that could interfere with cell biology or hamper medical use. It is known, that the concentration of signalling molecules decides the cell fate (schematical representation Figures 1D, F modified by (Huizar et al., 2020). Thus, the tip geometry and coating of the Sphericalplate 5D is designed to support this process. Sharp or flat tips in classical microwell platforms interfere with this process and do not allow equal fate decisions because of unequal signalling molecule concentrations (Figures 1E, F). Thus, the Sphericalplate 5D is fulfilling these prerequisites for future cell therapies. This is protected by IP.

### Establishment and characterization of pancreatic pseudoislets

In a first step, human pancreatic islets were plated either in commonly available cell culture dishes (fresh intact), multiwell plates (Intact MWP) or as re-aggregated pseudoislets on Sphericalplates (300 or 600 cells per pseudoislet). Afterwards, important basal characteristics such as Ki67 as proliferation marker, the islet area per cell number, pyknotic bodies and Cleaved-Caspase-3 as markers reflecting cell death, furthermore Glucagon, PPY, Somatostatin and Insulin were determines. Subsequent quantifications revealed overall constant parameters with the exception of basally increased insulin levels for pseudoislets consisting of 600 cells each. Thus, our experiments confirmed general applicability for high throughput establishment of pseudo-islets in round tip wells without any need for additional anti-adherence buffers.

### Establishment and characterization of adrenal tumor spheroids from adrenocortical cell lines

Next, for translation foor adrenal-related applications, the human adrenocortical cell lines NCI-H295R and MUC-1 were seeded to Sphericalplates and investigated for successful tumor spheroid formation as well as ongoing spheroid growth as exemplified in Figure 3 B and C. Histological and immunohistochemical stainings confirmed viable spheroids (Figure 3 A and D), which reflected tumor model specific proliferation rates and steroidogenic activities as demonstrated by both SF-1 and 3betaHSD positivity (Figure E and F).

**Figure 2:**
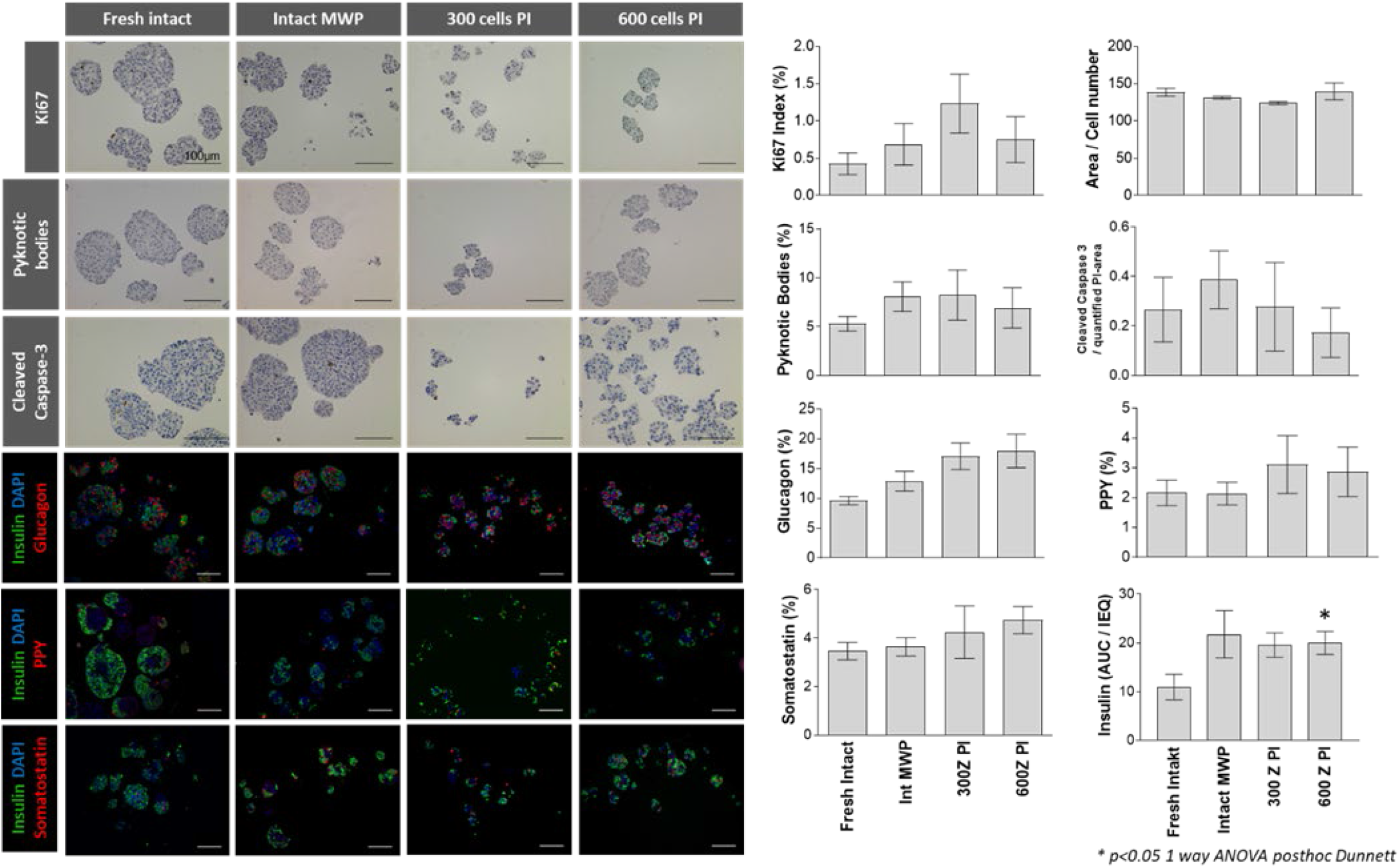
Freshly isolated islets were either cultured intact (fresh islets), in multiwell plates (Intakt MWP) or dissociated to form pseudoislets (PI) constituting of 300 cells (300 cells PI) or 600 cells (600 cells PI). Islets or pseudoislets from each condition were processed for IHC for various markers, scale bar 100um. Cells positive for the markers were counted and reported as percentage of all cells. Data obtained from 10 individuals (n=10; including furthermore quantifications from three high power fields per individual) is reported as mean +/- sem and was analyzed by ANOVA with Dunnett test *p<0.05.

**Figure 3:**
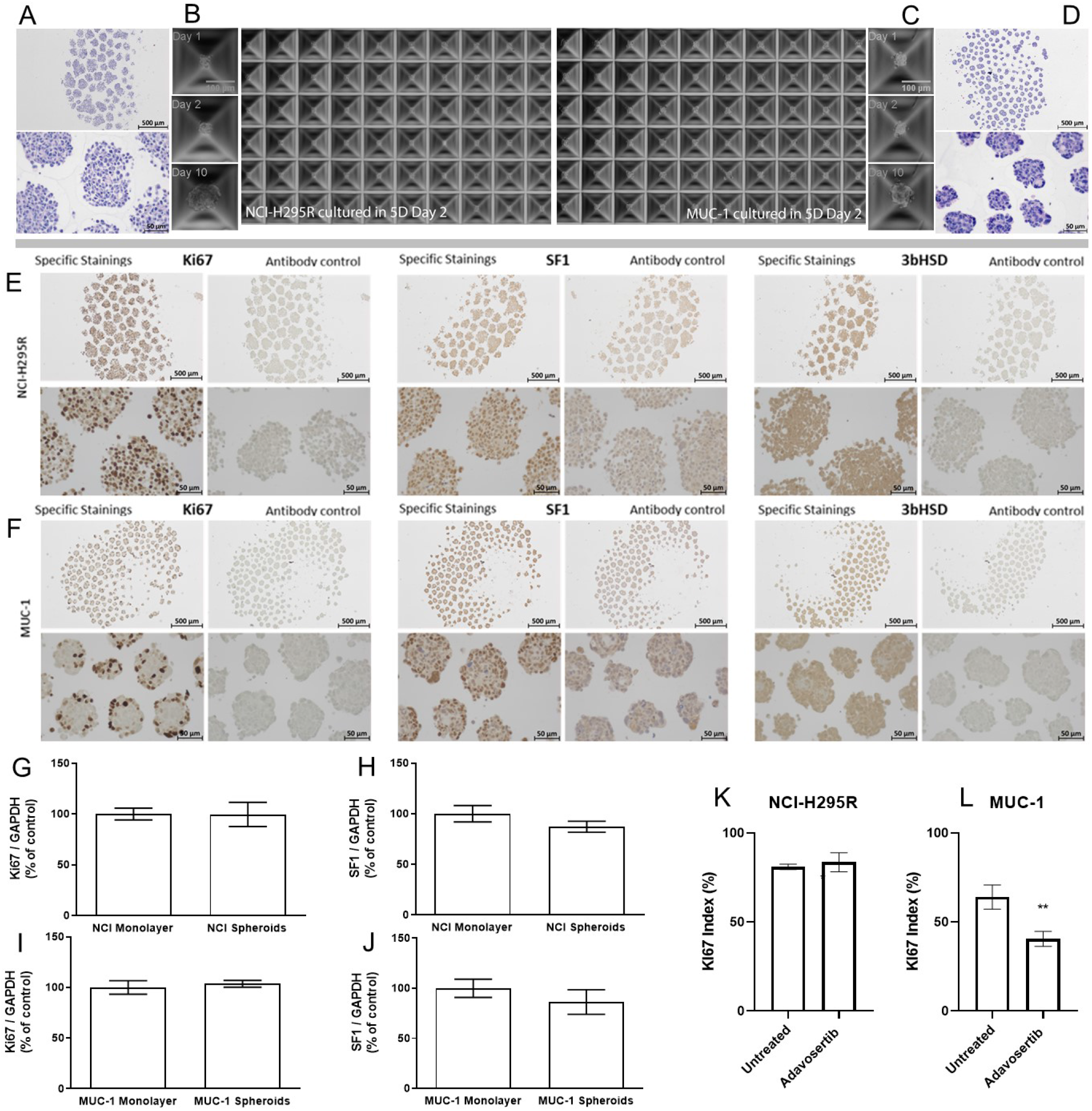
NCI-H295R and MUC-1 cell lines form spheroids in the Sphericalplate 5D. H&E staining of NCI-H295R spheroids (A) and MUC-1 spheroids (D) and progression of spheroid formation (B and C respectively) over 10 days. NCI-H295R spheroids (E) and MUC-1 spheroids (F) were processed for IHC with antibodies against Ki67, SF-1 and 3bHSD, scale bar 500μm and 50μm. Gene expression of Ki-67 (G) and SF-1 (H) was compared by real-time RT-PCR in NCI-H295R monolayer cells and spheroids (n=3). Gene expression of Ki67 (I) and SF-1 (J) was compared by real-time RT-PCR in MUC-1 monolayer cells and spheroids. Ki67 index between control and Adavosertib-treated NCI-H295R (K) and MUC-1 (L) spheroids. Data analyzed by Student’s t-test, **p<0.01.

Subsequent real time PCR investigations for potential changes in Ki-67 or SF-1 gene expression levels in tumor spheroid versus the appropriate monolayer cultures, did not reveal significant differences (Figure 3 G-J). Pilot experiments investigating the calculation of Ki-67 Index upon specific drug treatments as important pathological determinant, demonstrated furthermore applicability of such readouts. The quantification of Ki-67 indices revealed significant anti-proliferative effects of adavosertib, a small molecular Wee1-inhibitor, against MUC-1 but not NCI-H295R tumor spheroids. These results were in agreement with parallel MTT experiments on monolayer cultures (data not shown).

### Establishment and characterization of patient-derived tumor spheroids from adrenocortical origin

To extend the established adrenal tumor models for applications in terms of personalized medicine, we aimed furthermore at the development of patient-derived tumor spheroids. Figure 4A demonstrates appropriate H&E, Ki-67, SF-1 and 3betaHSD stainings which overall accounted for viable and proliferating spheroids derived from a primary, non-secreting ACC.

**Figure 4:**
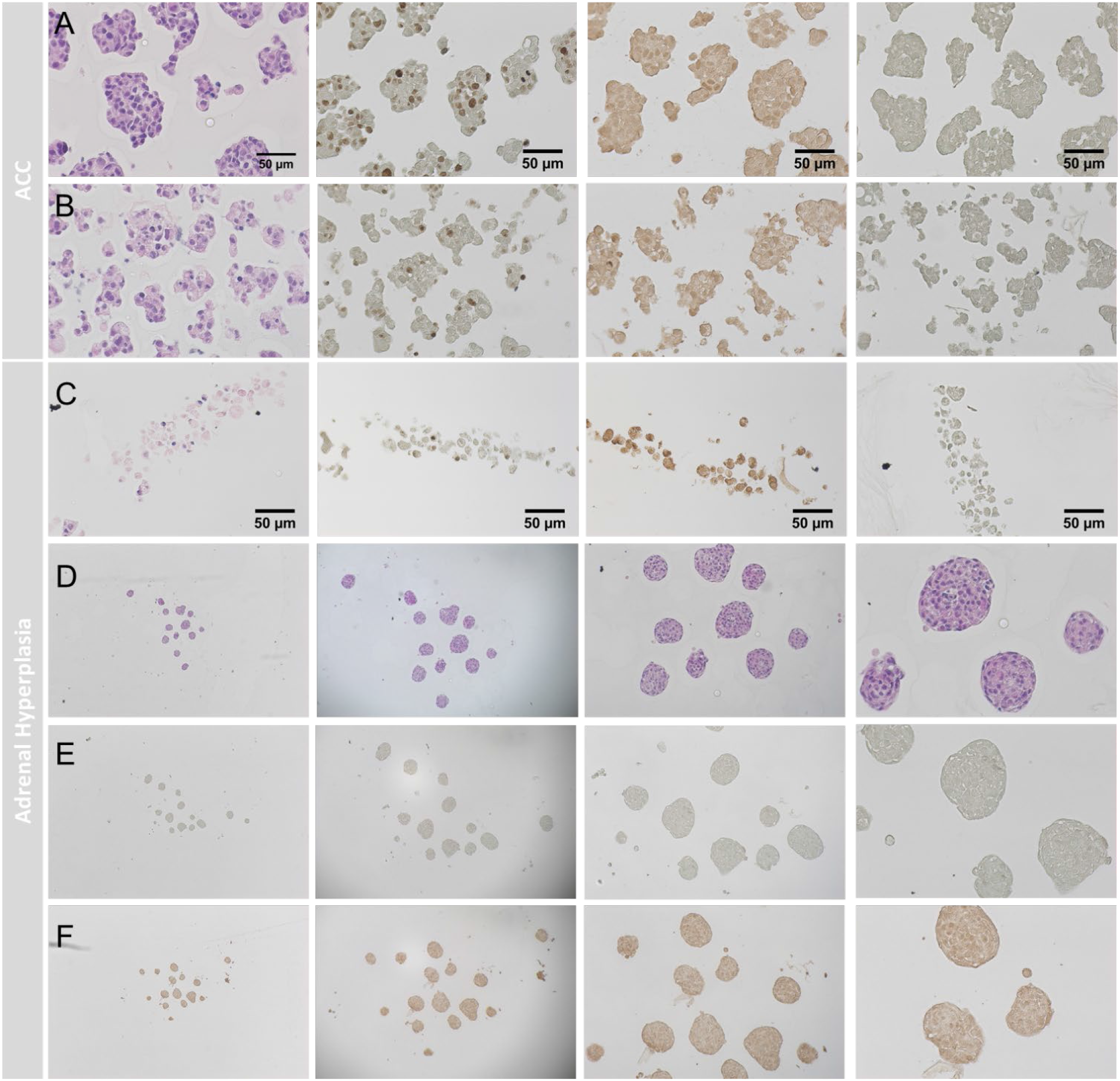
Spheroid formation from patient-derived ACC investigated by H&E, Ki-67, SF-1 and 3bHSD. Control spheroids (A), spheroids treated with 25μM Gemcitabine (B) and spheroids treated with Gemcitabine (25μM) and Cisplatin (40μM, C). Patient-derived spheroids from benign adrenal hyperplasia investigated by H&E (D), Ki-67 (E) and SF-1 (F).

Even though for SF-1 elevated background staining was detectable for these spheroids, the experiments indicated clearly definable nuclear SF-1 positivity compared to the appropriate antibody controls, but rather low or absent 3betaHSD abundance. Histologically, weak anti-tumoral effects were already detectable upon treatment with 25 μM Gemcitabine (Figure 4B). Figure 4C demonstrates severe pathological changes upon treatment with 25 μM Gemcitabine and 40 μM Cisplatin applying the same stainings as listed above. Interestingly, not only malignant tumors can be cultured by this method, Figure 4D shows viable spheroids of a benign adrenal hyperplasia, with the characteristic low/lacking Ki-67-Index (Figure 4E) compared to ACCs (Figure 4A). These tumor-spheroids were furthermore SF-1 positive (Figure 4F), but 3betaHSD negative (data not shown).

### Establishment and characterization of patient-derived tumor spheroids from medullary origin

Despite the need of innovative multi-dimensional and patient-derived *in vitro* models for adrenocortical tumors, there is nowadays also still a fundamental lack of models of medullary origin with no human cell line available. In an attempt to investigate putative applicability also for this tumor entity, we cultured cells freshly obtained from a surgical tumor sample of a benign pheochromocytoma. Of note, as demonstrated in Figure 5, we obtained viable tumor-spheroids (5A), which showed no proliferation *in vitro* (5B), but high functionality as indicated by strong chromogranin A stainings (Figure 5C vs. appropriate antibody controls 5D). Test-wise administration of 25 μM Gemcitabine, confirmed again applicability of such read-outs. However, patient-individual the treatment resulted for these spheroids in no therapeutic response as compared with the pictures above for ACC tumor spheroids.

**Figure 5:**
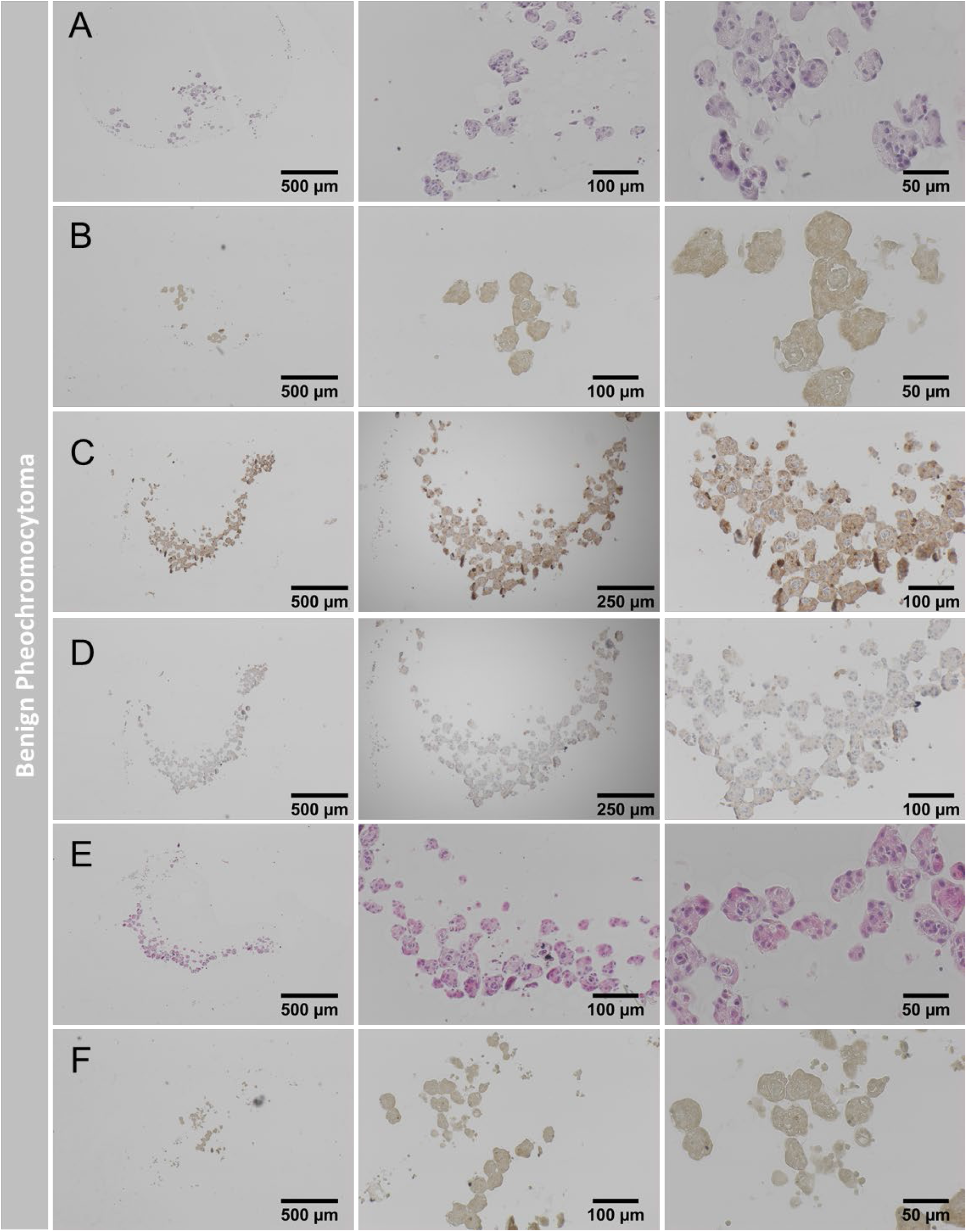
Spheroid formation from benign pheochromocytoma. H&E staining of spheroids at 3 different magnifications (A). Spheroids were analyzed with the proliferation marker Ki-67 (B). Spheroids processed for chromogranin A (C) and its respective no-antibody control (D). Spheroids treated with 25μm Gemcitabine (H&E, E and Ki-67, F).

### Establishment and characterization of organoids from normal adrenal origin (bovine and porcine)

Next, we aimed at the development of multi-dimensional models obtained from healthy adrenal glands (Figure 6).

**Figure 6:**
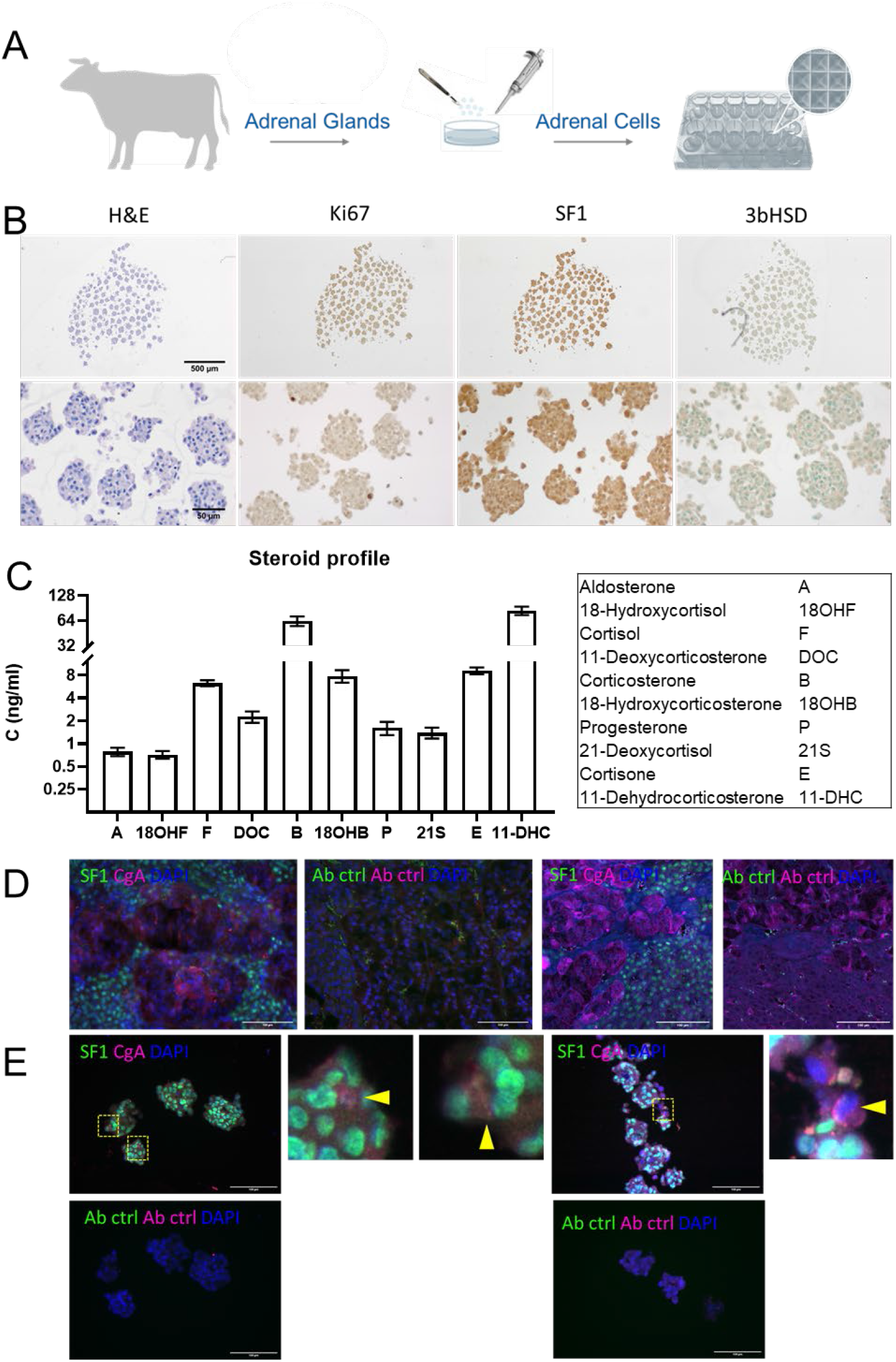
Successful formation of organoids from bovine cells of adrenocortical and medullary origin. Scheme depicting experimental set up (A). Organoids processed for H&E, Ki67, SF-1 and 3bHSD staining at two different magnifications. Secretory profile of organoids derived from a bovine adrenal gland as analyzed by LC-MS/MS (n=3) (C). Bovine adrenal sections stained with antibodies against SF-1 and chromogranin A (CgA), scale 100μm (D). Adrenocortical and medullary cells in organoids labeled with SF-1 and chromogranin A, scale 100μm (E).

For this purpose, we made cross-sections from healthy adrenals to reflect this time cells of both, adrenocortical and medullary origin, *in vitro* in one well. Bovine cells were subsequently cultured in Sphericalplates and as summarized in Figure 6B our experiments revealed viable and as for the healthy tissue origin expected low/not proliferating organoids. SF-1/3betaHSD immunohistochemistries and LC-MS/MS based adrenocortical steroid profiling confirmed specific basal steroidogenic activity (Figure 6B, C). Subsequent SF-1 and CgA immunefluorescence co-stainings verified furthermore viable co-cultures of cells specifically derived from both, adrenocortical and medullary origin (Figure 6D bovine adrenal tissue controls, 6E different organoids including antibody controls) indicating the potential for mimicking of dynamic zonation processes and subsequent investigation of zone to zone communication in vitro. In parallel, we currently investigate spheroids of bovine (Figure 7 A-C) vs. porcine (Figure 7 D-F) origin for potentially best applicability in terms of clinical transplantations for the treatment of diseases such as adrenal insufficiency. Both types demonstrate abundance of relevant markers such as DAX-1, Nestin and Sonic hedgehog indicating, thereby, stem cell and adrenal regenerative properties. Interestingly, culturing of porcine adrenal spheroids in NB-Medium led furthermore to highly improved cortisol and aldosterone output under basal and ACTH-stimulated conditions (Figure 7 G,H). Moreover, multicellular extension-formation could be observed under ACTH-treatment exclusively (Figure 7 J) compared to controls (Figure 7I). Of note, these types of multicellular filopodia demonstrated adrenal specific StAR, SF-1, CYP11A1 and CYP11B1 protein abundances (Figure 7 J-M). Electron microscopy showed furthermore, that these structures indeed originate from zona fasciculata cells (Figure 7 N).

**Figure 7:**
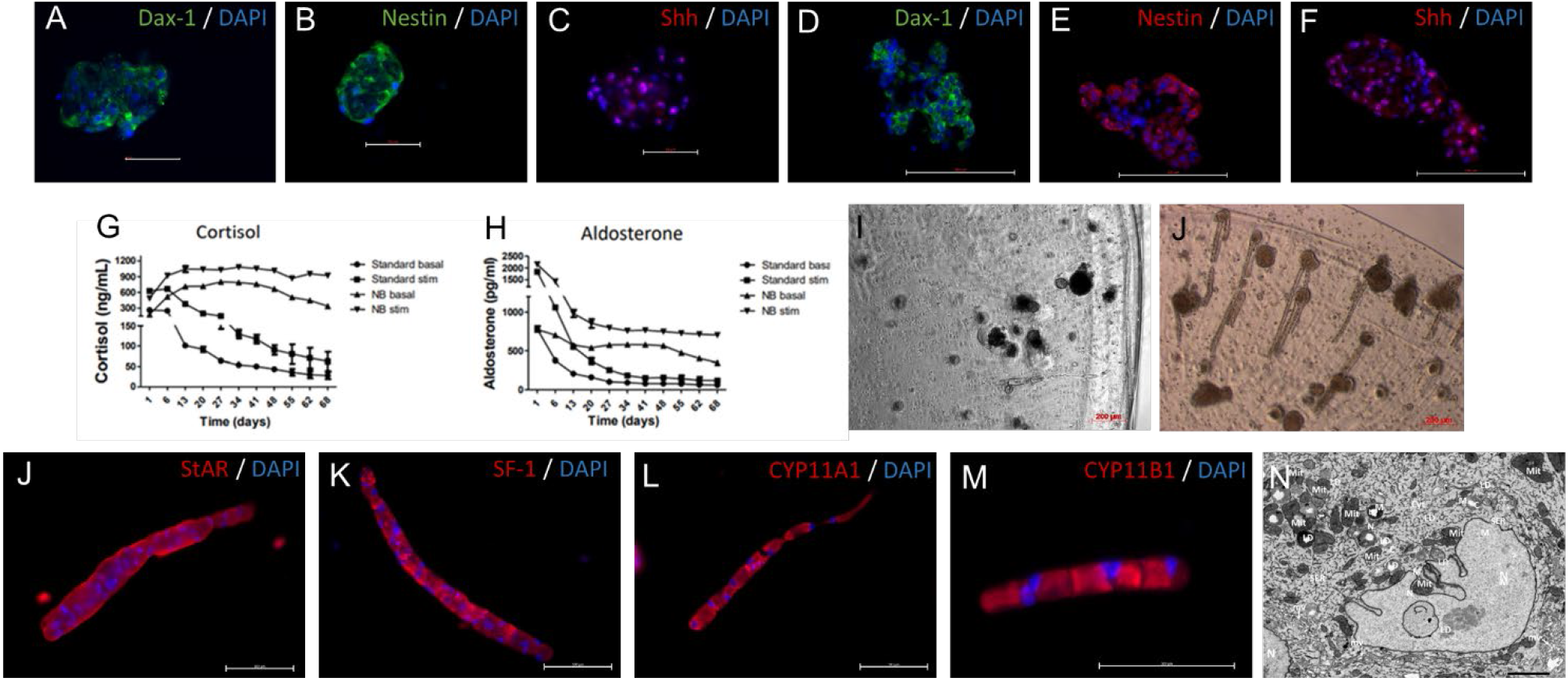
Spheroids from bovine (A-C) and porcine (D-F) adrenals stained with antibodies against DAX1, Nestin and SHH. Effect of growth media on the secretion of cortisol (G) or aldosterone (H) into the supernatant of porcine adrenal spheroids (3 experiments with 4 replicates each). Images of porcine spheroids (I) or spheroids treated with ATCH (J) highlight the presence of multicellular extensions following ACTH stimulation. Multicellular extensions expressed the markers StAR, SF-1 CYP11A1 and CYP11B1 (J-M). Electron micrograph of adrenocortical cells, migrating from the spheroid, containing adrenal progenitors, to the centre. The cell shows characteristic features of mature steroid producing cell including characteristic elongated and round-shape mitochondria (mit) with the typical tubulovesicular internal membranes. Furthermore, cells exhibit ample smooth endoplasmic reticulum (SER) and cell membrane extensions in form of microvilli (mv), scale 2 μm.

## Discussion

Here we report on the development of a multiwell plate created for multi-dimensional cell modelling and its applicability for different endocrine cell and tissue sources. As culturing of small pseudo-islet has been previously demonstrated to be advantegous over large islets (Lehmann et al., 2007) and is since then established in the field of diabetology, we focussed in our study on the confirmation of general applicability for such approaches. Moreover, we outlined general benefits such as round tip wells and no need for anti-adherent buffers. New potential applications for our subsequent described 3D-models in the adrenal field, however, range from basic research, cancer research for a wide variety of heterogeneous benign and malignant adrenal tumors to organ replacement therapies for adrenal insufficiency (de Witte et al., 2020; B. Pinto et al., 2020). Moreover, also chromaffin cells from the adrenal medulla have been considered e.g. as a potential source of dopamine-producing cells to treat neurodegenerative conditions like Parkinson disease or pain (Bornstein et al., 2020). First approaches for various tissue and species origins have been recently published. For example for organoids of specific human fetal adrenal origin general viability and expression of zonal and functional markers of the original fetal organ was demonstrated, but not for the implementation of post-natal tissues. Moreover, this technique and up to our knowledge all others applied until today, clearly lack high-throughput applicability (Poli et al., 2019). Our group previously demonstrated that alginated bovine adrenal spheroids can be a cell source for xenogeneic transplantation for the treatment of adrenocortical deficiency (Balyura et al., 2015). However, while originally developed to keep the cells isolated from the host immune response by a semipermeable membrane, it is known now that the immune system recognizes pathogen associated molecular patterns in alginate preparations themselves, thereby leading to adaptive and innate immune responses against engrafted capsules. Accordingly, the few clinical trials based on such alginate involving bioengineering approaches did finally not lead to any licensed therapy (Ashimova, Yegorov, Negmetzhanov, & Hortelano, 2019). The innovative spherical plate implemented in our novel approach, enables spheroid growth on a medical grade ultra-low attachment coating without encapsulation and even without additional need for anti-adherence rinsing solutions that could interfere with cell biology or hamper medical use. Moreover, there is substantial heterogeneity of various alginate capsules regarding the maximum permeability ranging from 25 to 250 kDa. These are additional factors which can also impact the normal exchange of spheroids regarding e.g. proteins and polysaccharides with their natural environment and thereby also viability (Ashimova et al., 2019). In contrast, the spherical plate enables a barrier-free exchange of spheroids and organoids with their appropriate environment and our data confirm viability as well as specific hormonal functionality. Of note, recently it has been shown that co-culturing of mouse or porcine cells together with CXCL12 leads to immunoprotection and long-term function with no further need of systemic host immunosuppression (Alagpulinsa et al., 2019). Consequently, we currently aim in parallel at investigations on porcine adrenal cells, their extra-ordinary regenerative potential and thereby optimized suitability for such approaches without or with low immune-activating stabilization in the future patient. Our most recent experiments demonstrate, while basal DAX1, Nestin and SHH abundances between bovine and porcine cells seemed to be comparable, the newly discovered ACTH-dependent potency to differentiate to StAR, SF-1, CYP11A1 and CYP11B1 consisting multicellular extensions indicate the extraordinary regenerative potential of porcine cell populations in this context. In next steps, we aim to combine and optimize our findings specifically for the establishment of high-throughput porcine derived adrenal spheroids immune-optimized for transplantation purposes.

Finally, our report demonstrates the most innovative and complete approach for 3D-modelling of adrenal tumors starting from two different adrenocortical tumor cell lines, benign, malignant as well as adrenocortical and medullary tumors. Up to date, except limited description of classically cultured NCI-H295R spheroids (Creemers et al., 2016; Haider, Schreiner, Kendl, Kroiss, & Luxenhofer, 2020; Lichtenauer et al., 2013), to our knowledge no comparable report, practical approach and appropriate protocol has been reported so far for adrenal tumors. Moreover, in addition to this overall proof of applicability for all described types of adrenal cells and tissues, it demonstrates furthermore superior properties over classical approaches as it also allows large-scale tumor spheroid formation, drug testings and readouts such as classical Ki-67 indices and thereby the potential applicability for personalized medical approaches in appropriate endocrine tumor boards.

## Materials and Methods

### Cell Culture

#### Human pancreatic islets

Human islets were obtained by the Islets for Research Distribution Programme of the European Consortium for Islet Transplantation (ECIT; biological replicates n=10, technical replicates n=3 from three high power fields per individual). The providing centres were the Cell Isolation and Transplantation Centre, Geneva University Hospitals; the Islet Processing Facility, S. Raffaele Scientific Institute Milan; “Recherche Translationnelle sur le Diabète” Faculté de Médecine, Université de Lille / CHRU de Lille / Inserm / EGID; and from Prodo Laboratories, Aliso Viejo CA. They were cultivated, dissociated into single cells and reaggregated as described previously (Zuellig et al., 2017). Dissociated cells were reaggregated in Sphericalplates 5D (Kugelmeiers, Erlenbach, Switzerland) at the required concentration in 0.5 ml medium (e.g. 300cells/microwell, for 750 microwells/well, 225000 cells/0.5ml). Medium was changed every second day. Islet perifusion was performed as described (Zuellig et al., 2017) except the device used was the Perifusion System (Model No: Peri-4.2, Biorep Technologies, Miami, FL).

#### Adrenocortical cell lines, primary culture and tumor spheroids

Primary cultures of patient tumors were established as previously reported (Hantel et al., 2016). Tumor sample collection was approved by the local ethics committee (Kantonale Ethikkommission Zürich, BASEC 2017-00771) and written informed consent was obtained from all subjects prior to tumor sampling. Cells from benign pheochromocytoma and benign adrenal hyperplasia were plated at 250000 cells /well in the spherical plate 5D (Kugelmeiers, Erlenbach, Switzerland). NCI-H295R, MUC-1 and ACC115m primary cells were cultured as previously described (Hantel et al., 2016; Rossini et al., 2021) and cultivated in Sphericalplates 5D for either 7 or 14 days at the densities of 75000 cells/well, 150000 cells/well and 75000 cells/well respectively. The 7 day spheroids were treated with 650nM Adavosertib (MedChemExpress) for 72 h. Additionally, ACC115m cells were cultivated in Sphericalplates 5D for 7 days and subsequently treated with 25μM Gemcitabine and 40μM Cisplatin (both Teva Pharma) for 72h.

#### Adrenal Spheroids

For 5D-cultured adrenal organoids and subsequent LC-MS/MS measurement of appropriate supernatants a 1.22g cross-section piece of bovine adrenal gland was minced to less than 0.5mm pieces using a razor blade in PBS. The resulting suspension was centrifuged at 250g for 5 minutes and the pellet was incubated with 2mg/ml sterile filtered Collagenase II (Gibco, Waltham) in DPBS for 50min at 37°C. Collagenase was inactivated with 5ml FBS (Thermo Fisher Scientific). Cells were pelleted and erythrocyte lysis was performed with 5ml RBC lysis buffer (pH 7.4) for 7min at room temperature (150mM ammonium chloride, 1mM potassium hydrogen carbonate and 0.1mM disodium-EDTA). Upon another centrifugation step, cell pellet was resuspended in PBS, and, after straining through a 70μm mesh, cells were counted using a Neubauer Improved hemocytometer, and centrifuged again at 250g for 5min. Dead cells were removed using Dead Cell Removal Kit (Miltenyi Biotec). Live cells were resuspended in Advanced DMEM/F-12 medium containing 17.51mM D-glucose, non-essential amino acids, and 1mM sodium pyruvate supplemented with 10% FBS and 1% penicillin/streptomycin (all Thermo Fisher Scientific), and seeded in Sphericalplates 5D at a density of 250000 cells/well.

For the investigation of stem cell and differentiation characteristics bovine adrenocortical cells were isolated from bovine adrenal glands shortly after the slaughtering of 1-3 years old cattle as previously described (Balyura et al., 2015). Briefly, adrenal glands were transported to the laboratory in ice-cold Euro Collins Solution supplemented with 1% (vol/vol) penicillin-streptomycin solution (Thermo Fisher Scientific). The glands were then liberated from fat and connective tissue and rinsed several times with PBS through the central vein to remove remaining blood. Afterwards a longitudinal incision was made to cut the adrenals in halves, the medulla was removed and the cortex was scraped off the capsule and cut in small pieces. Adrenal capsule was discarded. Adrenal cortex was digested for 50 min in DMEM/F12 medium (Thermo Fisher Scientific), containing 2 mg/ml collagenase and 0.1 mg/ml DNase (both from Sigma-Aldrich) at 37°C while shaking. Obtained cells were washed with cultivation medium, pelleted by centrifugation (8 min, 300 g) and filtered through 100-μm cell strainers (Becton Dickinson). After that, primary adrenocortical cells were placed in cell culture flasks (Thermo Fisher Scientific) and cultivated at 37°C in a humidified atmosphere (95% air, 5% CO_2_) in DMEM/F12 medium with 10% (vol/vol) FBS, 10% (vol/vol) horse serum (both from Thermo Fisher Scientific), 0.1 ng/ml recombinant FGF-2 (PromoCell GmbH) and 1% (vol/vol) penicillin-streptomycin solution. Medium change was performed every 2-3 days. Cells remained in culture for 4 days.

For spheroid formation, frozen cells were thawed, resuspended in standard medium and cultivated for 3 days. Cells were then trypsinized and placed in cell culture inserts (1 × 10^6^ cells per insert) with PTFE membrane (Merck) in 6 well plates for overnight cultivation. The next day the spheroids from each insert were mixed gently with 150 μl of 2% alginate (previsously described (Balyura et al., 2015), placed 30 μl of the mixture on a glass with 550 micron spacers, covered with flat Sintered glass (Pyrex) and cross-linked with 70 mM strontium chloride for 10 min. Received slabs were cultivated in neurobasal medium (Neurobasal medium, containing 2% (vol/vol) of B-27 supplement, 2 mM L-glutamine (all from Thermo Fisher Scientific), 20 ng/ml FGF −2 (Promocell) and 1 % (vol/vol) penicillin-streptomycin-nystatin).

#### Porcine adrenal spheroids

Porcine adrenal cells were isolated from adrenals of freshly slaughtered pigs. During the transportation and cell isolation, all solutions, except enzyme, contained 1% (vol/vol) of penicillin-streptomycin-nystatin (Neolab). The glands were transported to the laboratory in cold Euro-Collins solution, freed of connective and adipose tissue and washed twice in PBS. Then the glands were cut in half by a longitudinal incision, the cortex with medulla were scrapped off the capsule, cut into small pieces, and digested for 50 min in DMEM/F12 (Gibco, Thermo Fisher Scientific), containing 2 mg/ml collagenase type V (Sigma-Aldrich), and 0.1 mg/ml DNase (Roche) at 37°C while shaking. The cells were then washed twice in standard cultivation medium and pelleted by centrifugation (8 min, 300 x g) and filtered through gauze. Primary adrenal cells were cultivated in standard medium (DMEM/F12 with 5% (vol/vol) FBS, 10% (vol/vol) horse serum (both from Gibco, Thermo Fisher Scientific), 10 μg/ml insulin (Sanofi), 5 μg/ml transferrin (Sigma-Aldrich) and 1 % penicillin-streptomycin-nystatin) at 37 °C in a humidified atmosphere (95% air, 5% CO_2_). On the day following cell isolation, the cells were removed from the cell culture by trypsinization (TrypLE™, Thermo Fisher Scientific), pelleted by centrifugation, resuspeneded in a freeze medium (standard medium, containing 10% (vol/vol) DMSO) and stored frozen in the liquid nitrogen.

### Immunohistochemistry

#### Pancreatic pseudo-islets

Spheroids were fixed with 4% paraformaldehyde (PFA, Sigma Aldrich), embedded in Histogel (Thermo Fisher Scientific), then dehydrated and processed for paraffin embedding using the Excelsior AS tissue processor (Thermo Fisher Scientific). Spheroids were embedded using Myr embedding center (Especialidades Médicas MYR) and 4μm sections were prepared (microtome Leica RM2255, Leica Biosystems). For Ki67 staining, slides were processed with the Autostainer Link48 (DAKO, Agilent) and all reagents were purchased from DAKO. For all other antibodies, sections were deparaffinized, rehydrated, and, following the HIER treatment in sodium citrate buffer, incubated with blocking buffer containing 3 % BSA (Roche Diagnostics), 5 % goat serum (Jackson ImmunoResearch Laboratories), and 0.5 % Tween 20 (Sigma Aldrich). Sections were incubated with the following primary antibodies overnight: Ki67 (1:100, IR626, DAKO), cleaved caspase 3 (1:300, 9661, Cell Signaling), insulin (1:200, I2018Sigma-Aldrich), glucagon (1:50, PA5-13442, ThermoFisher), ppy (1:200, RP030-05, Diagnostic BioSystems) and somatostatin (1:250, A0566, DAKO). The following day, sectiones were either processed with the Vectastain Elte ABC-HRP kit (Vector Laboratories) and visualized by DAB (Sigma Aldrich) or detected with fluorescent secondary antibodies (Jackson ImmunoResearch). Images were processed with Zeiss Axiovert 200M with AxioCamMRc5 (6 images/slide).

#### Adrenal spheroids

Spheroids were processed as outlined in the pancreatic islet section. Primary antibodies used with the adrenal cortical cell lines were incubated overnight at 4C and were as follows: Ki67 (1:200, DCS), SF-1 (1:100, Perseus Proteomics) and 3β-HSD1 (1:1500, provided by Anita Payne, Stanford University School of Medicine, Stanford, CA, USA). Secondary antibodies were applied for 1h at room temperature (for Ki67 and 3βHSD1: 1:200, goat anti-rabbit biotinylated IgG, Vector Laboratories; for SF1: ImmPRESS™ HRP Anti-Mouse IgG (Peroxidase) Polymer Detection Kit, Vector Laboratories). For Ki67 and 3βHSD1, the Vectastain Elite ABC-HRP Kit (Vector Laboratories) was applied. Bound antibodies were visualized by 3,3’-diaminobenzidine staining (Sigma Aldrich).

Immunofluorescence stainings and imaging of spheroid sections were performed as described previously (Steenblock et al., 2021) using following antibodies: SF-1 (1:200, Cosmo Bio), Chromagranin A (1:100, Santa Cruz Biotechnology), donkey anti-goat-Cy3 (1:500, Jackson ImmunoResearch), and donkey anti-rabbit Alexa Fluor 647 (1:500, Invitrogen).

### Immunofluorescence of bovine and porcine adrenal spheroids

Free floating adrenal spheroids were fixed overnight in 4% PFA and embedded in Tissue-Tek O.C.T (Sakura Finetek). Immunohistochemistry was performed on 6-μm sections using primary antibodies for DAX1 (1:100, LS-B3480, LSbio), Nestin (1:100, NBP1-02419, Novusbio), SHH (1:100, LS-C40460, LSbio), Following overnight incubation with primary antibodies, sections were washed and incubated with for 1 h at room temperature with secondary antibodies goat anti-rabbit Alexa Flour 568 (1:1000, A11011; Life Technologies) for Nestin, SHH, or goat anti-rabbit Alexa Flour 488 (1:1000, A11034, Life Technologies) for DAX1, nestin or goat anti-mouse Alexa Flour 488 (1:1000, A11001, Life Technologies). DAPI (10236276001; Roche) was used for cell nucleus-specific staining. Immunofluorescence microscopy was performed on an Axioplan Carl Zeiss microscope, using Axiovision Software, or on Zeiss Axiovert 200M with AxioCamMRc5, using ZEN 3.3 (Carl Zeiss) software.

Spheroids encapsulated in alginate and adrenal column were washed with PBS and fixed with 4% PFA overnight. Since the fixative is hypo-osmotic, during the fixation procedure the alginate swells, which increases its porosity. After washing with PBS, the slabs were incubated in blocking solution (3% BSA, 0.1% Triton X-100 in PBS for 30 min at room temperature), then incubated with the primary antibodies DAX1, SHH, WNT4 (1:100, ab91226, Abcam), StAR (1:100, sc-25806, Santa Cruz Biotechnology), SF1, CYP11A1 (1:100, sc-18043, Santa Cruz Biotechnology), and CYP11B1 overnight at 4°C. The following day, slabs were washed with PBS and incubated for 1 h with the secondary antibody (goat anti-rabbit Cy3-conjugated IgG, 1:500, 111-165-144, Jackson Immunoresearch laboratories). Cell nuclei were stained with DAPI. Leftovers of alginate were removed by incubating in 500 mg/ml (w/v) solution of alginate lyase (Sigma-Aldrich) for 30 min. Immunofluorescence microscopy was performed on Axioplan microscope, using Axiovision Software.

### Transmission Electron Microscopy (TEM)

The spheroids were fixed in 4% PFA in 0.1 M phosphate buffer (PB, pH 7.4), followed by postfixation in modified Karnovsky’s fixative (2% glutaraldehyde + 2% paraformaldehyde in 50 mM HEPES, pH 7.4) overnight at 4°C (Juznic et al., 2021; Kurth et al., 2010). Samples were washed 2x in 100 mM PB and 2x in water and postfixed in 2% aqueous OsO4 solution containing 1.5% potassium ferrocyanide and 2 mM CaCl2 for 30 min on ice. After washes in water, the samples were incubated in 1% thiocarbohydrazide in water (20 min at room temperature), followed by washes in water and a second osmium contrasting step in 2% OsO4/water (30 min, on ice). Samples were washed in water, en bloc contrasted with 1% uranyl acetate/water for 2 h on ice, washed again in water, dehydrated in a graded series of ethanol/water (30%, 50%, 70%, 90%, 96%, 3x 100% ethanol (pure ethanol on molecular sieve)), and infiltrated in the epon substitute EMBed 812 (epon/ethanol mixtures: 1:3, 1:1, 3:1 for 1.5 h each, pure resin overnight, pure resin 5 h). Finally, the samples were embedded in flat embedding molds and cured at 65°C overnight. Ultrathin sections were prepared with a Leica UC6 ultramicrotome (Leica Microsystems, Vienna, Austria), collected on formvar-coated slot grids, and stained with lead citrate (Venable and Coggeshall, 1965) and uranyl acetate. Contrasted ultrathin sections were analysed on a Jeol JEM1400 Plus (JEOL, Freising, Germany, camera: Ruby, JEOL) running with 80 kV acceleration voltage.

### Steroid and insulin measurements (LC-MS/MS and ELISA)

For functional assessment of steroid hormone release by adrenal spheroids liquid chromatography tandem mass spectrometry (LC-MS/MS, technical replicates n=3) was applied as described elsewhere (Peitzsch et al., 2015).

For basal and ACTH stimulated steroid release, the slabs were incubated with or without 3 ng/ml ACTH_1-24_ (Synachthen, Sigma-tau Arzneimittel GmbH) for 24 h. Measurements of cortisol, aldosterone or insulin in cell culture supernatants were done by cortisol, aldosterone (IBL) and insulin (mercodia Cat. Nr. 10-1113-01) ELISA, respectively.

### Real-time PCR

Spheroid and monolayer samples (technical replicates n=3) of MUC-1 and NCI-H295R cells were processed for RNA extraction using the RNeasy Mini kit (Qiagen), followed by DNA removal (TURBO DNA-free™ Kit, Thermo Fisher Scientific) and reverse transcription (RevertAid™ H Minus First Strand cDNA Synthesis Kit, Thermo Fisher Scientific) where 310 ng of RNA was converted to cDNA per sample. For Real-Time PCR analyses, we utilized the SsoFast EvaGreen® reaction mix (Bio-Rad Laboratories) in the 7500fast cycler (Applied Biosystems). The following PCR primers were used: human SF1 (forward: 5’-CAGCCTGGATTTGAAGTTCCT, reverse: 5’-CAGCATTTCGATGAGCAGGT) and human Ki67 (forward: 5’-TCCTTTGGTGGGCACCTAAGACCTG, reverse: 5’-TGATGGTTGAGGCTGTTCCTTGATG). Gene expression levels were normalized to the housekeeping gene GAPDH (forward: 5’-AGCCTCCCGCTTCGCTCTCT, reverse: 5’-CCAGGCGCCCAATACGACCA).

### Statistical Analysis

Statistical analysis and graphical representation of the data was carried out using GraphPad Prism software (version 8, GraphPad Software, La Jolla, CA, USA). Statistical comparisons were performed using one-way ANOVA followed by Bonferroni or Dunnett’s multiple comparisons test. All results are expressed as mean ± SEM. The statistical significance was defined as P < 0.05 and denoted as stars in the graphs (* p < 0.05; ** p< 0.01; *** p< 0.001; **** p< 0.0001) in all figures, if not stated otherwise.

## Acknowledgements

We would like to thank Alessia Scapin for her excellent technical assistance with the characterization of the human pseudoislets.

